# Structure–activity relationship study in soil identifies an effective broad-spectrum inhibitor of archaeal nitrification

**DOI:** 10.64898/2026.10.29.762912

**Authors:** Laure Annetta, Fabian Beeckman, Kenzo Vereecken, Tom Beeckman, Hans Motte

## Abstract

Ammonia-oxidizing archaea (AOA) can contribute substantially to nitrification in agricultural soils, yet the portfolio of AOA-targeting nitrification inhibitors (NIs) with demonstrated efficacy under soil conditions remains limited. Moreover, the performance of candidate inhibitors in soil is often difficult to predict from pure-culture assays because their activity is strongly influenced by soil-dependent factors. In this study, we employed a recently developed high-throughput miniaturized soil incubation assay to perform a soil-based structure-activity relationship (SAR) analysis of SIAS (Specific Inhibitor of AOA in Soil), a pyrazolo-pyridine derivative previously identified as an AOA-targeting compound with strong additive nitrification inhibition effects when combined with the AOB-targeting inhibitor DMP (3,4-dimethyl-1H-pyrazole). To identify structural features associated with improved and more robust soil efficacy, a library of 27 SIAS-derived pyrazolo-pyridine analogues was evaluated directly in soil. The SAR analysis showed that inhibitory activity was governed by a specific combination of structural features and led to the discovery of Optimized SIAS (Opt. SIAS), a simplified analogue that consistently outperformed the original lead compound across diverse agricultural soils. Correlation analysis further revealed that soil pH and, particularly, silt content were major determinants of Opt. SIAS efficacy. The strong additive effects obtained through combined application of Opt. SIAS and DMP highlight the contribution of AOA to residual nitrification following AOB inhibition and demonstrate the potential of simultaneously targeting both groups. Together, these results expand the limited portfolio of effective AOA-targeting nitrification inhibitors and provide a foundation for the development of next-generation nitrification inhibition strategies aimed at improving nitrogen-use efficiency and reducing nitrogen losses from agricultural soils.

## 1. INTRODUCTION

Nitrogen (N) is an essential macronutrient required for optimal plant growth and is therefore applied extensively in agricultural systems to sustain high crop yields and meet the increasing global food demand (FAO, 2025; Godfray et al., 2010). However, considerable amounts of nitrogen fertilizers applied to agricultural soils serve as substrates for microbial nitrification, a process that contributes substantially to nitrogen losses from agricultural systems, resulting in nitrogen-use efficiencies often below 50% (Bowles et al., 2018; Norton and Ouyang, 2019). When N is supplied predominantly as ammonium (NH_4_^+^) or urea, rapidly hydrolysed to NH_4_^+^ by soil urease, it stimulates the microbial nitrification process (Cantarella et al., 2018; Huang et al., 2019). During nitrification, ammonia (NH_3_), which is in a pH-dependent equilibrium with NH_4_^+^ is oxidized to nitrite (NO_2_^−^) and subsequently to nitrate (NO_3_^−^) (Beeckman et al., 2018). This transformation is mediated either in two steps by Gram-negative ammonia-oxidizing bacteria (AOB) and ammonia-oxidizing archaea (AOA), or in a single step by complete ammonia-oxidizing (comammox) bacteria (Daims et al., 2015; Hayatsu et al., 2021; van Kessel et al., 2015). As such, nitrification leads to nitrogen losses from agricultural soils through NO□□ leaching, leading to groundwater contamination, or through gaseous emissions such as nitric oxide (NO) and nitrous oxide (N_2_O) produced during anaerobic denitrification, nitrifier denitrification or biotic/abiotic conversion of metabolic intermediates (Sutton et al., 2011).

To mitigate eutrophication and greenhouse gas emissions while maintaining high crop productivity, nitrification inhibitors (NIs) have been applied alongside N fertilizers for several decades (Lam et al., 2022). Currently approved NIs for agriculture, like 3,4-dimethyl-1H-pyrazole (DMP) and dicyandiamide (DCD), primarily suppress ammonia oxidation by AOB (Papadopoulou et al., 2020; Zhou et al., 2020). Nitrapyrin is stated to target both AOB and AOA, however, it does not clearly affect AOA abundance in soil (Fisk et al., 2015; Hayden et al., 2021; Shen et al., 2013). Furthermore, the total nitrification inhibition can still be enhanced by combining nitrapyrin with an AOA-targeting inhibitor (Beeckman et al., 2023), indicating a predominant AOB-targeting as well. Although AOB are generally considered the dominant ammonia oxidizers in agricultural soils, AOA may contribute more to nitrification under certain conditions (Beeckman et al., 2024), e.g. in acidic soils (Gubry-Rangin et al., 2010), or when AOB growth is inhibited by AOB-targeting NIs (Zhou et al., 2020). These observations highlight the potential of specifically targeting AOA in addition to AOB for improved nitrification inhibition. In laboratory settings, several AOA-targeting NIs have been identified, including PTIO (2-phenyl-4,4,5,5-tetramethylimidazoline-1-oxyl-3-oxide) and ethoxyquin – presumably acting as NO scavengers (Papadopoulou et al., 2020; Shen et al., 2013) – as well as simvastatin, which disrupts archaeal cell integrity (Zhao et al., 2020). However, the efficacy of these compounds under soil conditions has been investigated only to a limited extent and, in some cases, their specificity toward AOA remains debated, as reported for PTIO (Fu et al., 2018; Lin et al., 2023).

Recently, the AOA-targeting compound 4-[1,6-bis(propan-2-yl)-1H-pyrazolo[3,4-b]pyridine-4-carbonyl]thiomorpholine (SIAS – for Specific Inhibitor of AOA in Soil), was identified through a culture-based screening assay using the AOA *Nitrososphaera viennensis*. SIAS was subsequently shown to improve nitrification inhibition in soil when combined with nitrapyrin or AOB-targeting inhibitors such as DMP, presumably due to the simultaneous targeting of both AOB and AOA (Beeckman et al., 2023), and corroborating the significant contribution of AOA to nitrification under certain conditions. However, not all SIAS-like structures that effectively inhibited the AOA *Nitrososphaera viennensis* showed this additive effect in soil, highlighting the limited ability of culture-based assays to predict inhibitor performance under soil conditions (Beeckman et al., 2023), likely because sorption, mobility, and bioavailability strongly influence the activity of agrochemicals in soil (Strawn, 2021). Therefore, identifying AOA-targeting inhibitors that remain effective across diverse soil conditions when used in combination with an AOB-targeting NI, and understanding which structural features govern their performance, are important steps toward developing improved nitrification inhibition strategies.

In this study, to identify the structural determinants of effective AOA inhibition in soil and optimize SIAS-like structures, we performed a structure–activity relationship (SAR) analysis of SIAS-derived pyrazolo–pyridine compounds directly in soil, and investigated how these determinants influence inhibitor performance across a broad range of agricultural soils.

## 2. MATERIALS & METHODS

### 2.1. Miniaturized soil nitrification inhibition assay

The miniaturized soil nitrification inhibition assay was performed according to Antoniou et al. (2026). In brief, the agricultural soils collected in Belgium and Germany or ordered at Lufa-Speyer (https://www.lufa-speyer.de/en) (**Table 1**), were air-dried and sieved (mesh diam. 1 mm), and transferred to each well of a 96 deep-well plate (AB-0661, Thermo Fischer scientific), using a 96-well PCR plate (Cat. No. 732-2387, VWR) as a mold. A watery solution containing glucose monohydrate (1.289 g kg^−1^ dry soil) and ammonium nitrate (0.003 g kg^−1^ dry soil) was added to all wells to activate microbial C- and N-cycles. The volume of activation solution was adapted for each soil in order to reach a water filled pore space (WFPS) of 55%. After a short centrifugation (5 s), the plates were sealed off with sealing mats (AB-0675, Thermo Fisher Scientific) and incubated in the dark at room temperature (20 °C) for 14 days. Then, 5 μL of either a watery solution of 10% dimethyl sulfoxide (DMSO, Sigma Aldrich) as mock control, or a watery solution of 1 mM NIs in 10% DMSO: 3,4-dimethylpyrazole (DMP, EuroChem), or 4-[1,6-bis(propan-2-yl)-1H-pyrazolo[3,4-b]pyridine-4-carbonyl]thiomorpholine (SIAS, Enamine), as positive controls (final dose of 0.005 µmol well^−1^ in soil). When combined with DMP, SIAS or SIAS variants (**Table 2**; **Table S1**) were mixed in equal proportions (at 1 mM) in 10% DMSO solution and both added in soil at a final dose of 0.005 µmol well^−1^. Additionally, 5 μL of 100 mM ammonium sulphate ((NH_4_)_2_SO_4_, Sigma Aldrich) solution was added to all wells (excl. the wells dedicated for the NH_4_^+^ standard curve), to reach a final dose of 1 µmol well^−1^ of ammonium (NH_4_^+^) in soil. Subsequently, the plates were centrifugated (5 s) then closed and incubated, in the dark at room temperature until read-out.

**Table 1:**
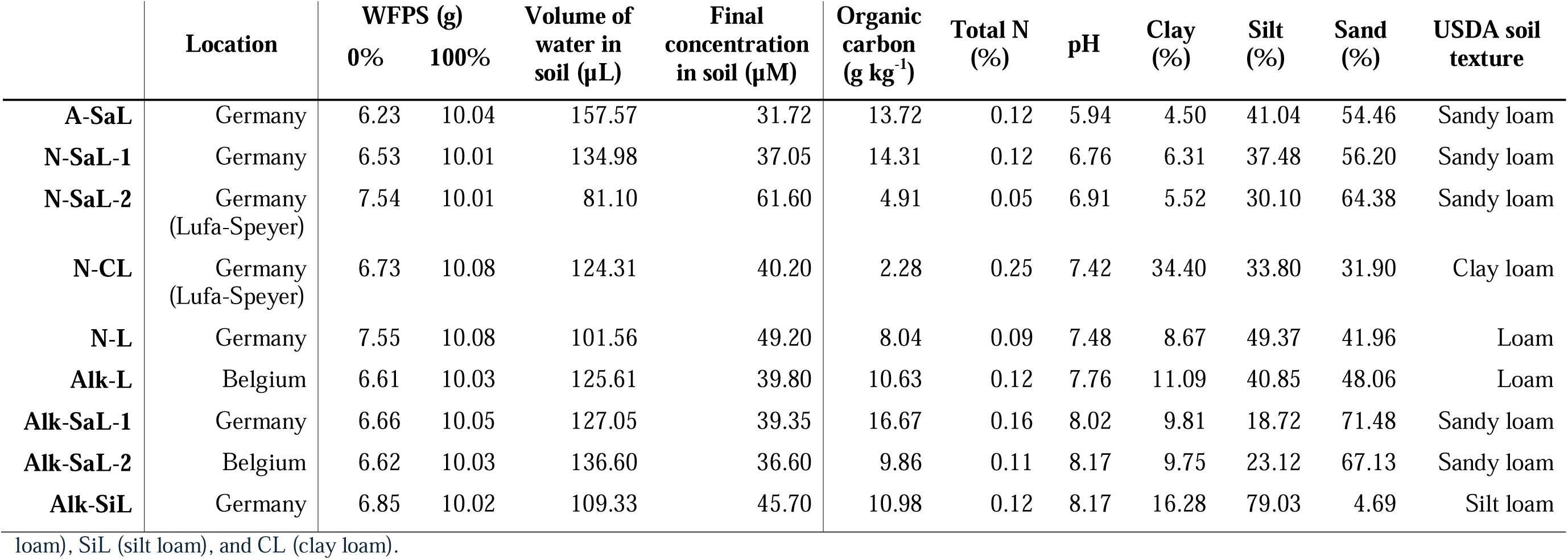
Chemical and physical properties of the studied soils, including organic carbon (g kg-1), total nitrogen, N (%), pH, particle size distribution in 128 % (clay, silt, and sand), 129 and USDA soil texture class. Total volume of water (μL) required to achieve a water-filled pore space (WFPS) of 55% calculated based on grams of soil corresponding to 0% 130 and 100% WFPS for each soil. Final compound concentrations are expressed in μM based on the volume of water contained in each soil. Soils are identified using a short 131 code based on pH class and USDA texture, where A = acidic (pH < 6.5), N = neutral (6.5–7.5), and Alk = alkaline (pH > 7.5); texture abbreviations are L (loam), SaL (sand loam), SiL (silt loam), and CL (clay loam).

**Table 2:**
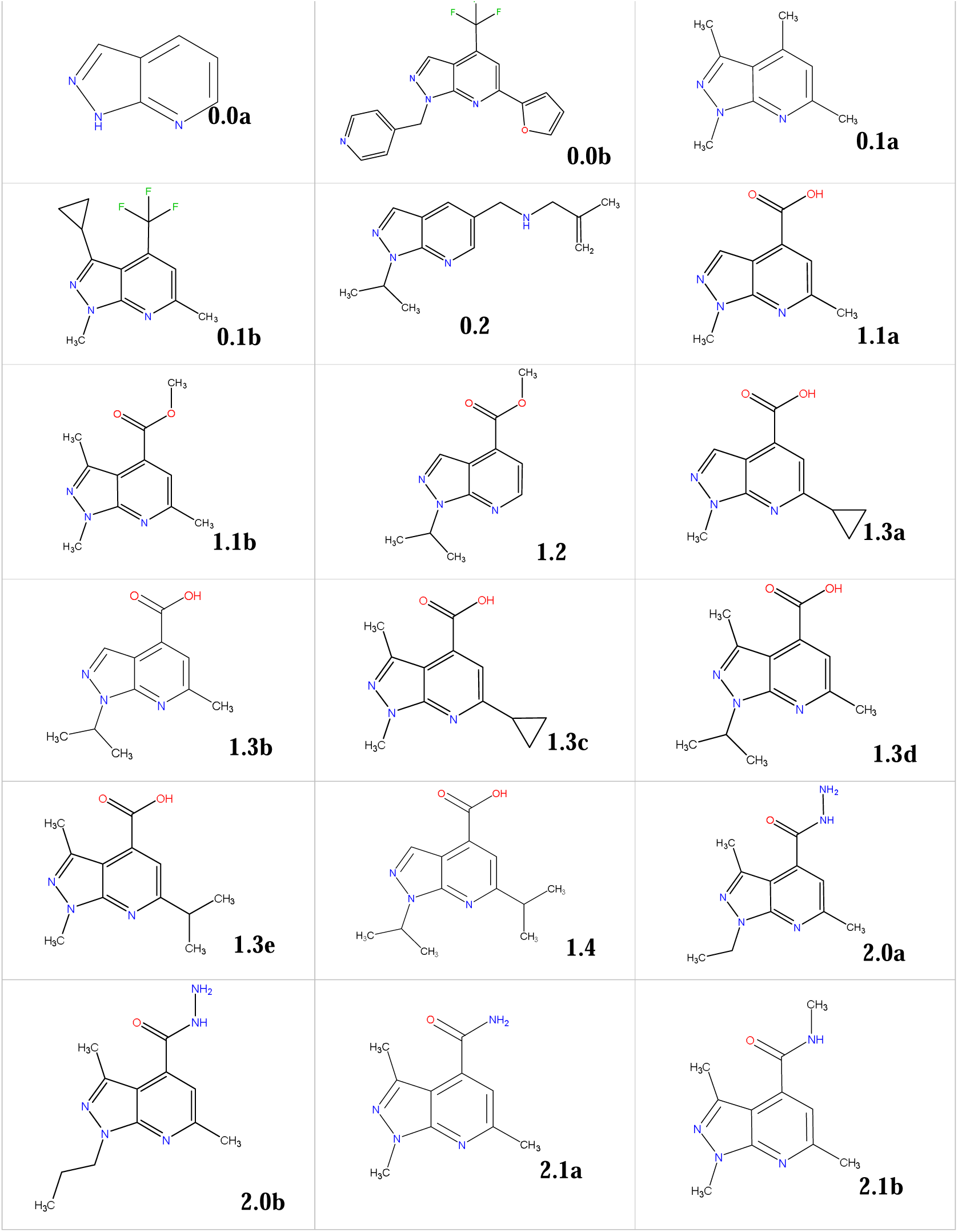
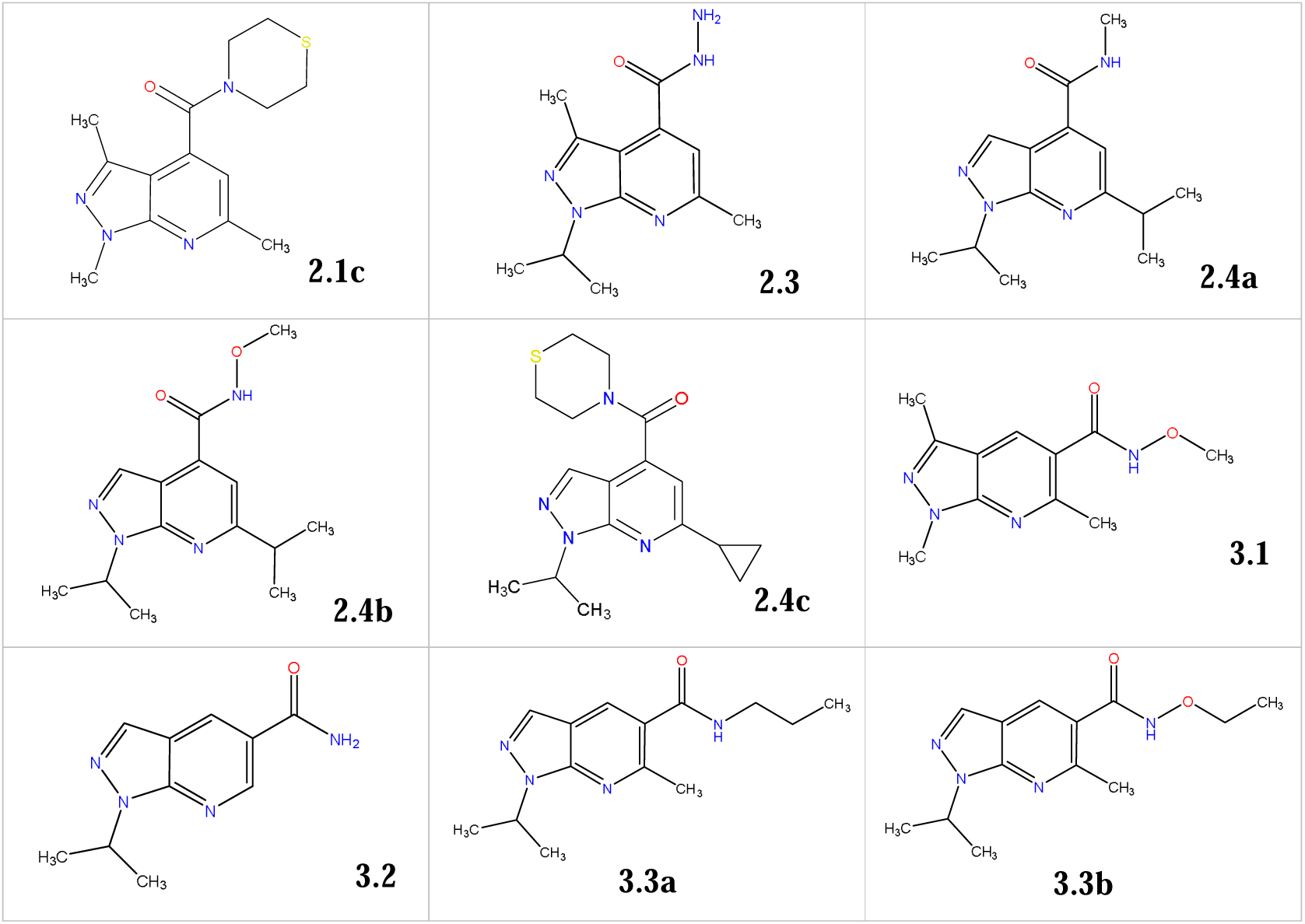
List of the 27 SIAS variants used in the structure-activity relationship (SAR). A detailed explanation of the compound numbering system is described in Figure 2.

At the end timepoint, 6-point serial dilutions of (NH_4_)_2_SO_4_ were added to the wells dedicated for the standard curve. Next, .1 mL of KCl (1 M, pH2) was added to the all the wells to stop nitrification and extract NH_4_^+^. Then, the plates were first shaken with a Retsch® machine (3 min, 20 Hz), on an orbital shaker (1 h, 500 rpm), and later centrifuged (10 min, 1000 rpm). NH_4_^+^ concentrations were measured using a modified Berthelot’s reagent protocol (Bucur et al., 2006) in a 96-well flat-bottom plate (Tissue culture plate, 96 wells-F, VWR®). Therefore, 168 μL of diluted sample was mixed with 35 μL of reagent A (125 mM sodium hydroxide and 2.4-2.6% sodium hypochlorite in Mili-Q water), and 33 μL of reagent B (72.4 mM salicylic acid, 125 mM sodium hydroxide, and 34.4 mM sodium nitroprusside dihydrate in Mili-Q water). After 30 min of incubation in the dark (room temperature), absorbances were measured at 635 nm using of a spectrophotometer (Versamax® Microplate reader). NH_4_^+^ concentrations (mM) were calculated using the standard curve.

### 2.2. *Nitrososphaera viennensis* nitrification inhibition assay

The cultures of *Nitrososphaera viennensis* EN76^T^ were grown and the nitrification assays were performed as described in Beeckman et al. (2023). In brief, *N. viennensis* was grown aerobically at 42 °C in fresh water medium (FWM) supplemented with 3 mM NH_4_Cl and 300 μM Na-pyruvate, according to Tourna et al. (2011), until late-log phase ([NO_2_^−^] ≥ 1 mM). The cell pellets of these cultures were then dissolved in fresh FWM and 5-times overconcentrated. Wells of a 96-well flat-bottom plates (VWR Tissue culture plate, 96 wells-F) were then filled with 198 µL of these overconcentrated cultures. Subsequently, 2 µL of the treatments were added to the wells, including: (i) 0.25% DMSO (mock control), (ii) PTIO (2-phenyl-4,4,5,5-tetramethylimidazoline-1-oxyl 3-oxide) at 10mM in 25% DMSO solution (positive control), or (iii) SIAS variant 2.4a/Opt. SIAS (**Table 2**; **Table S1**) at 2.5 mM in 25% DMSO solution (n = 6). Final concentrations in the culture were thus: 100 µM for PTIO or 25 µM for Opt. SIAS. Plates were incubated for 24h at 42 °C, in the dark. Nitrite (NO_2_^−^) measurements were performed using the modified Griess reagent (Sigma Aldrich) in equal proportions with (diluted) samples, in a 96-well flat-bottom plates (VWR Tissue culture plate, 96 wells-F). Later, absorbance values were evaluated with a spectrophotometer at 540 nm (Versamax® Microplate reader), and NO_2_^−^concentrations were calculated thanks to a standard curve.

### 2.3. Statistical analysis

Statistical analysis, Pearson correlations and simple linear regressions were achieved using GraphPad Prism (version 11.0.0).

## 3. RESULTS

### 3.1. The synergistic nitrification inhibition effect of SIAS and DMP is soil-dependent

The combined application of the AOA inhibitor SIAS (**Figure 1A**) with the AOB inhibitor DMP has previously been reported to enhance nitrification inhibition in two alkaline sandy-loam soils and one neutral loam soil (Beeckman et al., 2023). To further assess the robustness of this synergistic effect, we evaluated the impact of SIAS and DMP, alone and in combination, in a broader set of soils differing in texture and pH, including sandy-loam, loam, silt-loam, alkaline, neutral, and acidic soils (**Table 1**). To facilitate this large-scale evaluation, a recently developed miniaturized soil incubation assay was employed, providing substantially higher throughput than conventional soil incubation approaches (Antoniou et al., 2026). In these assays, a relatively low ammonium concentration was applied to enable fast consumption and facilitate the assessment of nitrification inhibition over short incubation periods. First, we characterized the rate of NH_4_^+^ consumption and the nitrification inhibition by DMP over time in each soil (**Figure 1B-F**). Based on these profiles, a time-point was selected at which NH_4_^+^ was (nearly) completely depleted in the mock treatment while DMP still exerted a partial inhibitory effect, enabling optimal discrimination between treatments. At this timepoint, NH_4_^+^ levels were quantified following treatment with SIAS, DMP, or their combination across all soils (**Figure 1G**). SIAS alone showed limited inhibitory activity in most soils, although it strongly inhibited nitrification on its own in the neutral loam soil (N-L). DMP strongly delayed ammonium consumption in most soils. Notably, the combined application of SIAS and DMP resulted in significantly higher NH_4_^+^ retention compared to DMP alone in the two alkaline soils, an effect that was confirmed in a full time-course experiment in the Alk-Sal soil (**Figure 1H**). In contrast, no additive effect was observed in the two neutral soils (N-SaL-1 and N-L) and the acidic soil (A-SaL) used in this experiment, indicating that the efficacy of the SIAS-DMP combination was strongly influenced by soil properties. Overall, SIAS did not act as an effective additive nitrification inhibitor across all soil types.

**Figure 1:**
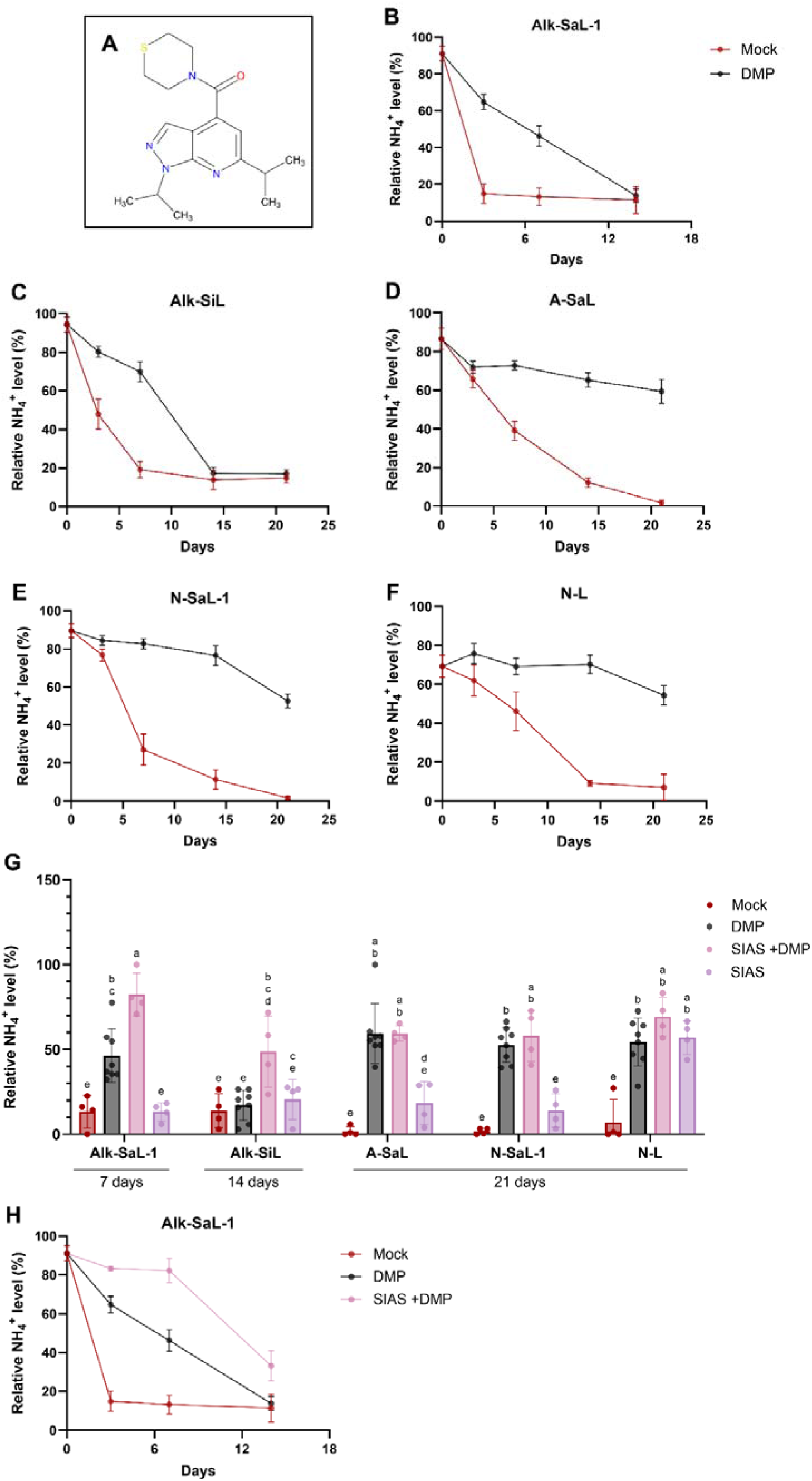
(**A**) Chemical structure of SIAS. (**B-F**) Relative ammonium (NH_4_^+^) level (%) in: Alk-SaL-1 (**B**), Alk-SiL (**C**), A-SaL (**D**), N-SaL-1 (**E**) and N-L (**F**) soils treated with DMP, or SIAS+DMP, evaluated at 0, 3, 7, 14 days (and 21 days for C-F). (**G**) Relative NH_4_^+^ level (%) in Alk-SaL-1 (at 7 days), Alk-SiL (at 14 days), A-SaL, N-SaL-1 and N-L (at 21 days) soils treated with DMP, SIAS or SIAS+DMP. Statistical analysis were assessed via a two-way ANOVA test followed by a Dunnett post-hoc test. (**H**) Relative ammonium (NH_4_^+^) level (mM) in Alk-SaL-1 soil treated with DMP, or SIAS+DMP, and evaluated at 0, 3, 7 and 14 days Treatments with different letters are significantly different (p<0.05). Errors bars represent SD (G) or SEM (B-F; H).

### 3.2. Structure-activity relationship analysis identifies structural determinants of soil efficacy

The soil-dependent performance of SIAS suggests that specific structural features may govern its efficacy under different soil conditions. To identify these features and explore opportunities for structural optimization, we assembled a library of 27 pyrazolo-pyridine derivatives sharing the SIAS core scaffold (**Table 2** and **Table S1**; **Figure 2**). All compounds were evaluated in an alkaline sandy-loam agricultural soil (**Table 1**) using the miniaturized soil incubation assay.

**Figure 2:**
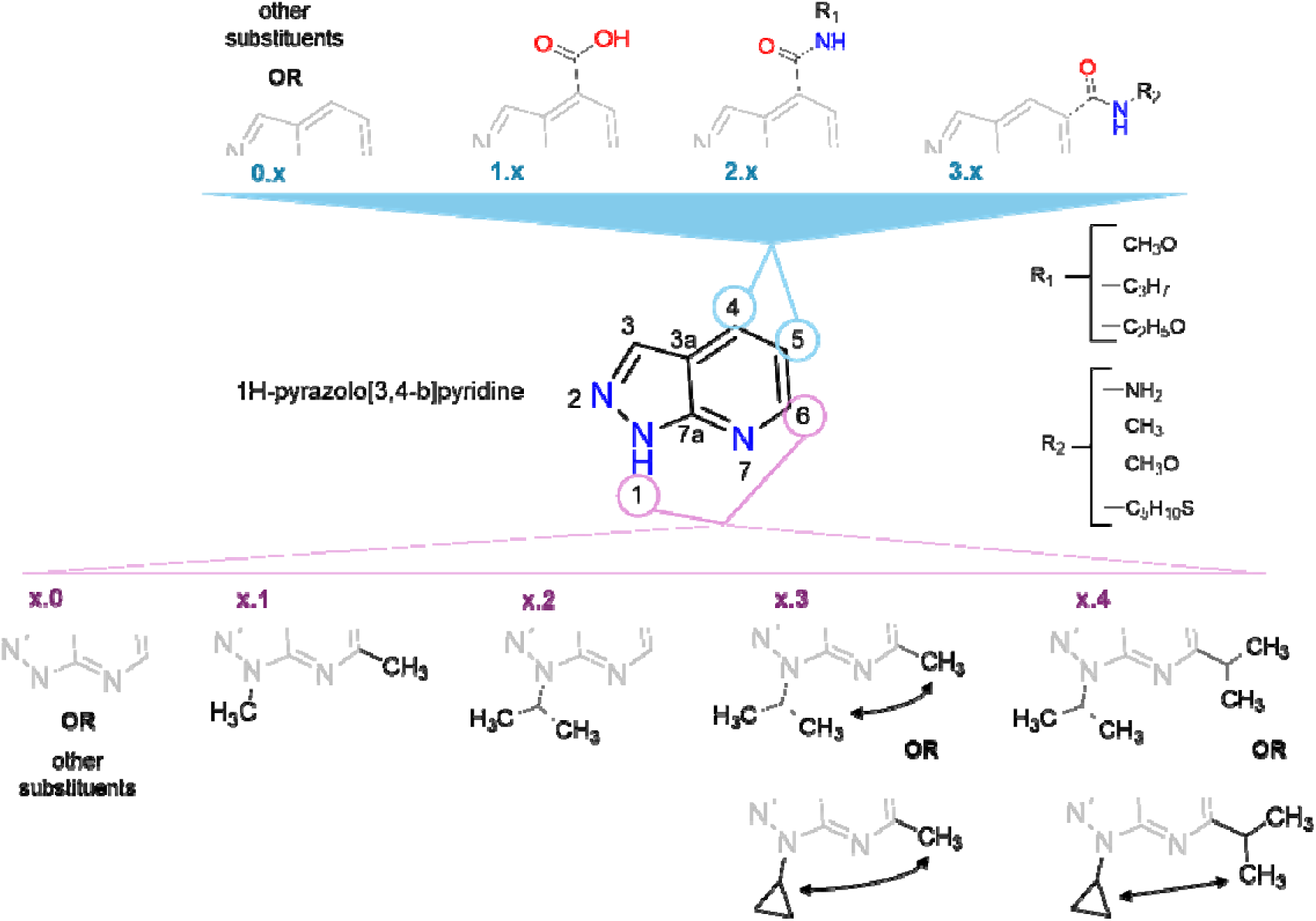
Principal structural variations present in the SIAS chemical variants used in this study. The compound designation consists of two numbers. The first number (0.x to 3.x) refers to the substitution at positions 4 or 5 of the pyrazolo-pyridine core: 0.x, none or other substituents than carboxylic acid/carboxamide substitution; 1.x, carboxylic acid (or carboxylate) at position 4; 2.x, carboxamide at position 4; 3.x, carboxamide at position 5. The second number (x.0 to x.4) refers to the substitution at positions 1 and/or 6: x.0, none or other substituents; x.1, two methyl groups; x.2, one isopropyl group; x.3, one isopropyl (or methyl-cyclopropane) and one methyl group; x.4, two isopropyl groups. R_1_ = – NH_2_, – CH_3_, – CH_3_O or – C_5_H_10_S. R_2_ = –CH_3_O, – C_3_H_7_ or – C_2_H_5_O.

Not all SIAS variants showed enhanced nitrification inhibition when applied in combination with DMP. In particular, most compounds from group 0.x, which lack carboxylic acid or carboxamide substitutions at position 4, did not result in significantly higher ammonium levels compared to DMP alone (**Figure 3A** and **C**). This indicates that the pyrazolo-pyridine core scaffold by itself is insufficient to confer additional inhibitory activity in soil. In contrast, the introduction of carboxyl-derived substituents at position 4 (groups 1.x and 2.x) generally, though not universally, increased ammonium retention, suggesting improved nitrification inhibition. However, compounds bearing similar substituents at position 4 did not consistently exhibit similar activity, indicating that additional structural features modulate activity. This is exemplified by compounds 2.1a and 2.1b: while 2.1a (**Figure 3A**), bearing a primary carboxamide, did not significantly enhance nitrification inhibition compared to DMP alone, the closely related analogue 2.1b (**Figure 3B**), containing an N-methyl-substituted secondary carboxamide, exhibited a clear additive effect. Similarly, variants containing an acetohydrazide substituent at position 4 (e.g. 2.0a and 2.0b) also demonstrated improved activity (**Figure 3C**). Notably, compound 2.4a, which carries the same N-methylcarboxamide group as 2.1b, showed an even stronger effect, exceeding that of SIAS, while 2.4b displayed a similarly enhanced activity (**Figure 3B**). These observations indicate that substituents at other positions on the pyrazolo-pyridine core significantly modulate inhibitory efficacy. Further evidence for the importance of substituent interplay is provided by modifications at position 5. Certain variants containing a carboxamide group at this position (e.g. 3.1 and 3.3a) exhibited significant nitrification inhibition, although generally weaker than SIAS (**Figure 3B** and **C**). In contrast, other compounds with similar substitutions (e.g. 0.2 and 3.3b) did not show additional activity (**Figure 3A** and **C**), further highlighting that nitrification inhibition depends on the combined structural context rather than on single functional groups alone.

**Figure 3:**
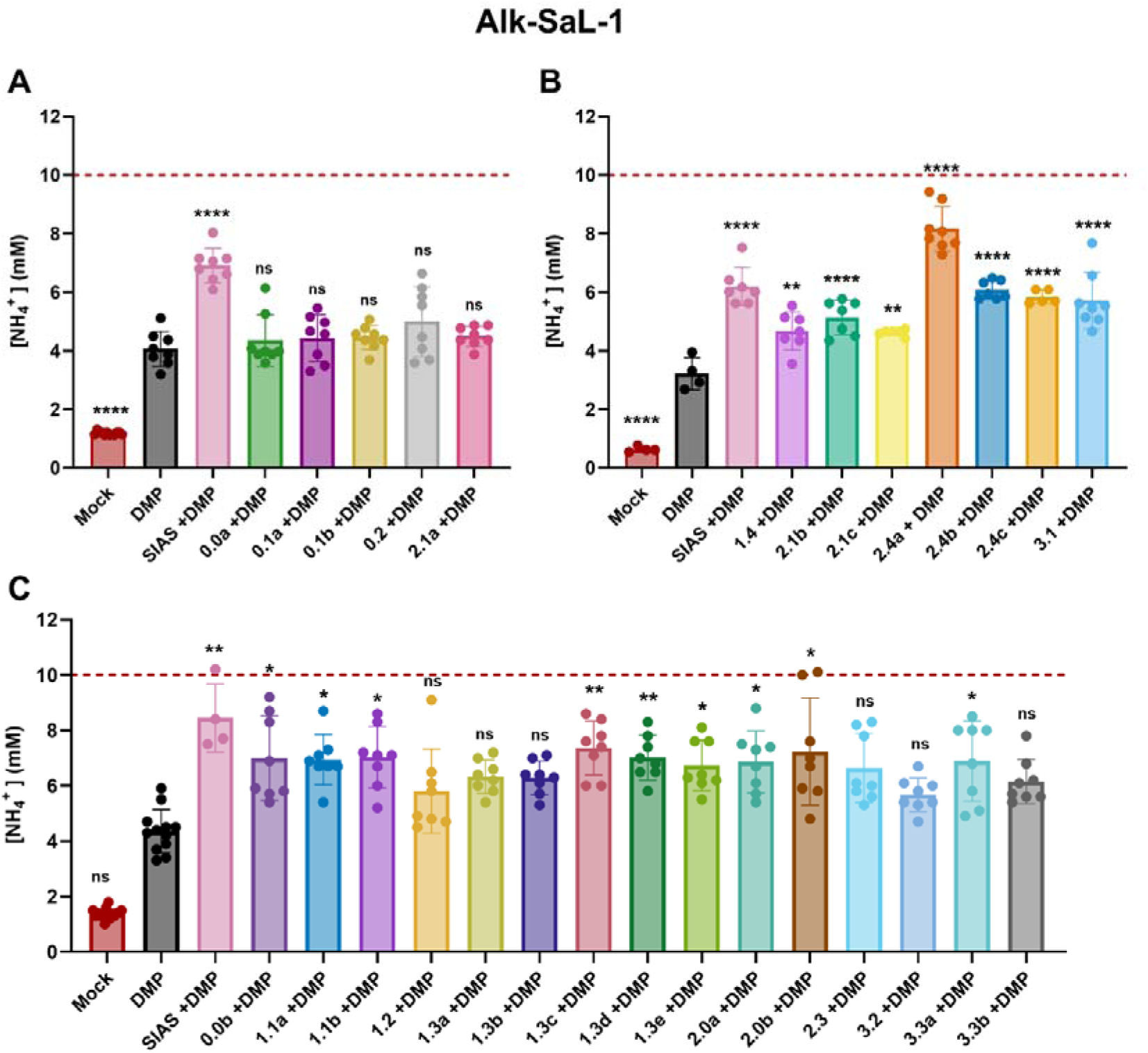
(**A-C**) Ammonium (NH_4_^+^) concentration (mM) in Alk-SaL-1 soil treated with DMP, SIAS or SIAS variants (numbers), in combination with DMP, and incubated for 7 days. Dotted line represents initial NH_4_^+^ level. Significant differences were assessed via a one-way ANOVA followed by a Dunnett’s test (A and B), or a Kruskal-Wallis test followed by a Dunn’s test (C). Treatments with an asterisk are significantly different from DMP (*: p<0.05; **: p<0.01; ***: p<0.001; ****: p<0.0001). Errors bars represent SD.

Variant 2.1c shares a similar overall structure with SIAS, containing a thiomorpholine substituent on the ester moiety; however, it lacks the two isopropyl groups at positions 1 and 6, which are replaced by methyl groups. This structural modification resulted in a reduced additive effect compared to SIAS when combined with DMP (**Figure 3B**). In contrast, variant 2.4c, which contains one isopropyl and one methyl-cyclopropane substituent at these positions, exhibited an activity comparable to SIAS, suggesting that large hydrophobic substituents at positions 1 and 6 contribute to enhanced inhibitory efficacy (**Figure 3B**). This trend is not absolute, as variant 1.4, which retains two isopropyl groups but carries a carboxylic acid at position 4, showed a reduced activity similar to 2.1c. This indicates that the beneficial effect of isopropyl substituents depends on the broader structural context. Additional modifications on the pyridine ring further influenced activity. One of the compounds bearing a trifluoromethyl group at position 4, variant 0.0b (**Figure 3C**), displayed moderate but significant increases in ammonium retention. By contrast, other substitutions at positions 1 or 6 (e.g. variants 0.0b and 2.0b) did not markedly alter activity, with these compounds showing effects comparable to SIAS.

Overall, the presence of a carboxamide or carboxylic group at position 4 or 5 of the pyrazolo-pyridine core (variants 1.x, 2.x and 3.x) appears to be necessary, but not sufficient, for soil activity. The observed activity furthermore depends on the nature of the substituents at the remaining positions. Especially a carboxamide on position 4 and iso-propyl substitutions on positions 1 and 6 positively affect the efficacy (variants 2.4).

To assess whether these structure–activity trends are conserved in another soil, a subset of representative compounds was further evaluated in a second alkaline soil (Alk-L -**Figure 4**). Here, most variants exhibited NH_4_ ^+^ levels comparable to DMP alone, indicating their limited contribution to nitrification inhibition under these conditions. Several compounds that displayed activity in the alkaline sandy-loam soil Alk-SaL-1, such as 1.4 and 2.1a, did not exhibit enhanced inhibition in this Alk-L soil, indicating a degree of soil-dependent variability. Nevertheless, compound 2.4a showed a strong and consistent performance across both soils. This further supports that specific substitution pattern, particularly those combining a carboxamide at position 4 with hydrophobic substituents at positions 1 and 6, are key determinants of enhanced nitrification inhibition in soil.

**Figure 4:**
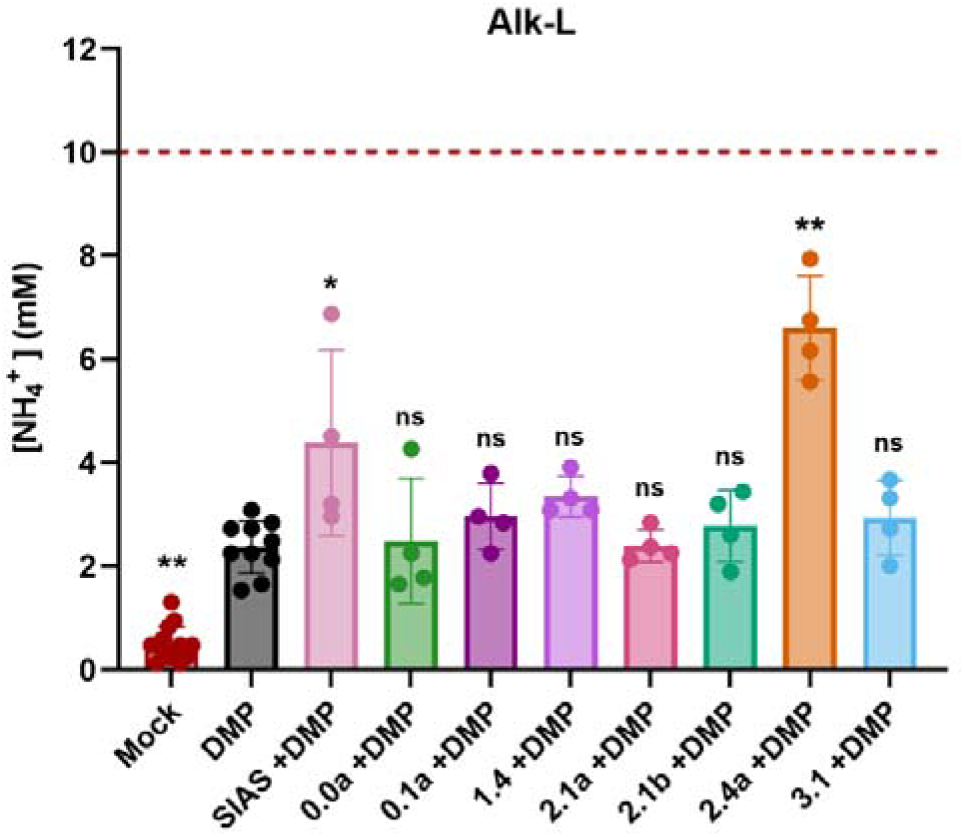
Ammonium (NH_4_^+^) concentration (mM) in Alk-L soil at 14 days treated with DMP, SIAS or SIAS variants, in combination with DMP. Dotted line represents initial NH_4_^+^ level. Statistical analysis was assessed via Kruskal-Wallis test, followed by an uncorrected Dunn’s test. Treatments with an asterisk are significantly different from DMP (*: p<0.05; **: p<0.01; ***: p<0.001; ****: p<0.0001). Errors bars represent SD.

### 3.3. Validation of structure–activity relationships across different agricultural soils

To further validate these findings and assess soil-dependent variability, a subset of seven representative variants was selected and tested in three additional sandy-loam agricultural soils, comprising one alkaline and two neutral soils (**Table 1**).

Across these soils, variants from the 2.4 series consistently exhibited a strong additive effect when combined with DMP (**Figure 5A–C**), confirming the initial SAR analysis. Likewise, the presence of a carboxamide group alone was not sufficient to ensure high activity. For example, variants 2.1b and 2.1c, both bearing a carboxamide group at position 4, exhibited only moderate or no significant enhancement compared to DMP alone, depending on the soil. In addition to the nature of the functional group, its position on the core scaffold proved to be critical. Variant 3.1, which carries a carboxamide at position 5 instead of position 4, and showing a significant additive effect in the Alk-Sal-1 soil, did not show significant activity in any of the other soils, indicating that a substitution at position 4 is generally more favourable for robust activity across soils. Similarly, compounds bearing a carboxylic acid at position 4 (e.g. variant 1.4) displayed only moderate effects, generally weaker than those observed for carboxamide-containing analogues. Notably, the relative performance of variant 1.4 varied between soils, further illustrating the influence of the overall structural context. A key determinant of activity appears to be the combination of a carboxamide group at position 4 with hydrophobic substituents at positions 1 and 6. Variants such as 2.1c, which lack the two isopropyl groups at these positions, showed reduced activity across soils. In contrast, compounds from the 2.4 series, which incorporate both a position 4 carboxamide and bulky hydrophobic substituents at positions 1 and 6, consistently exhibited the highest levels of ammonium retention. In particular, variants 2.4a and 2.4b demonstrated strong and statistically significant additive effects compared to DMP in all soils tested.

**Figure 5:**
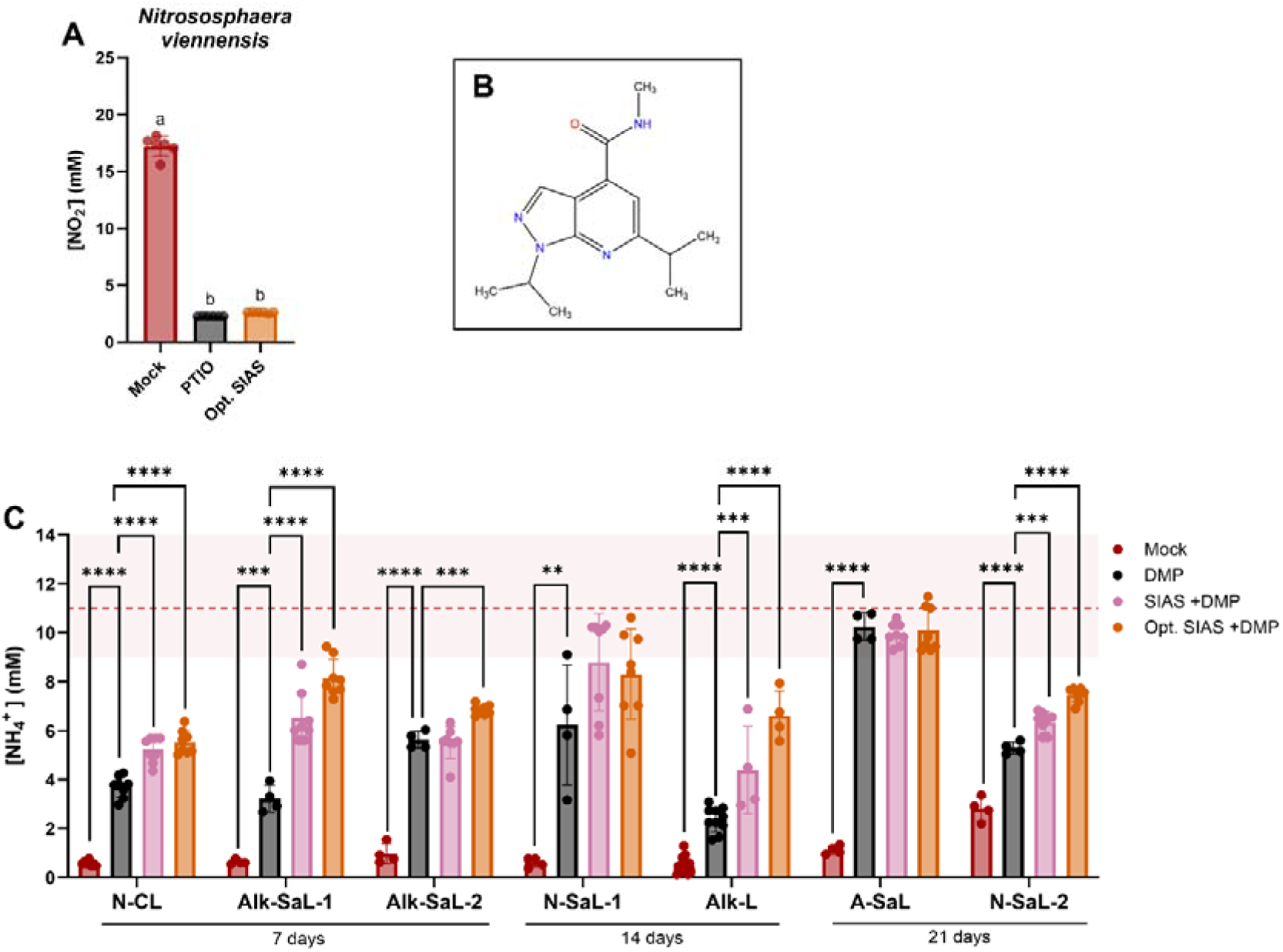
**(A)** Nitrite (NO_2_^−^) levels (mM) in pure cultures of the AOA, *Nitrososphaera viennensis*, treated with PTIO (100µM) or variant 2.4a or “Optimized SIAS” (Opt. SIAS) at 25µM. Statistical analysis was assessed via a one-way ANOVA test, followed by a Tukey’s test. Treatments with different letters are significantly different (p<0.05). **(B)** Chemical structure of Opt. SIAS. **(C)** Ammonium (NH_4_^+^) concentration (mM) in 7 soils: N-CL, Alk-SaL-1 and Alk-SaL-2 (at 7 days), N-SaL-1 and Alk-L (at 14 days), A-SaL and N-SaL-2 (at 21 days) treated with DMP, SIAS or Opt. SIAS, in combination with DMP. Dotted line represents average initial NH_4_^+^ level of the soils, and light red color, min and max. Statistical analysis was assessed via a one-way ANOVA test for each soil, followed by a Dunnett’s test. Treatments with an asterisk are significantly different from DMP (*: p<0.05; **: p<0.01; ***: p<0.001; ****: p<0.0001). Errors bars represent SD.

Taken together, these results validate the structure–activity relationship identified earlier (**Figure 3**) and demonstrate that optimal activity arises from a specific combination of structural features rather than from individual functional groups alone. Importantly, the consistent performance of 2.4 variants across multiple soils indicates that this substitution pattern confers a robust and largely soil-independent enhancement of nitrification inhibition. Among these, variant 2.4a consistently showed the strongest performance across diverse soil conditions, while having a simpler structure and lower molecular weight than the initial lead compound. In addition, it still strongly inhibited nitrite production by *Nitrososphaera viennensis* EN76 in pure culture, confirming that optimization of the SIAS scaffold did not compromise its AOA inhibitory activity (**Figure 5A**). This compound was selected for further investigation and hereafter referred to as “optimized SIAS” (Opt. SIAS) (**Figure 5B**). **Figure 5C** summarizes all soil incubation data specifically for SIAS and Opt. SIAS and includes additional data for the soils N-CL, Alk-SaL-2, N-SaL-1, A-SaL and N-SaL-2. It clearly shows that Opt. SIAS has an additive effect over DMP, besides in N-SaL-1 and A-SaL, and that the combined effect of Opt. SIAS and DMP consistently matches or exceed the effect observed with SIAS (**Figure 5C**).

### 3.4. Silt content and pH are key drivers of Opt. SIAS efficacy

To identify the soil parameters underlying the variability in the performance of Opt. SIAS, the additive effect of Opt. SIAS in combination with DMP was quantified and correlated with selected physicochemical soil properties using Pearson correlation analysis (**Figure 6A**). Among the parameters tested, only two showed a significant relationship with the additive effect of Opt. SIAS: silt content and soil pH. A strong negative correlation was observed with silt content, indicating that higher silt levels are associated with reduced efficacy of Opt. SIAS. In contrast, a positive correlation with soil pH suggests that the compound performs better in more alkaline conditions. These relationships were further evaluated using linear regression models (**Figure 6B-C**). The correlation with pH was moderate (R² = 0.47), indicating that approximately half of the variability in the additive effect can be explained by soil pH alone. In contrast, the relationship with silt content was substantially stronger (R² = 0.74), identifying silt content as a main determinant of Opt. SIAS performance across soils. Together, these results suggest that both soil texture and pH influence the efficacy of Opt. SIAS, with silt content representing the dominant factor governing its activity under the tested conditions.

**Figure 6:**
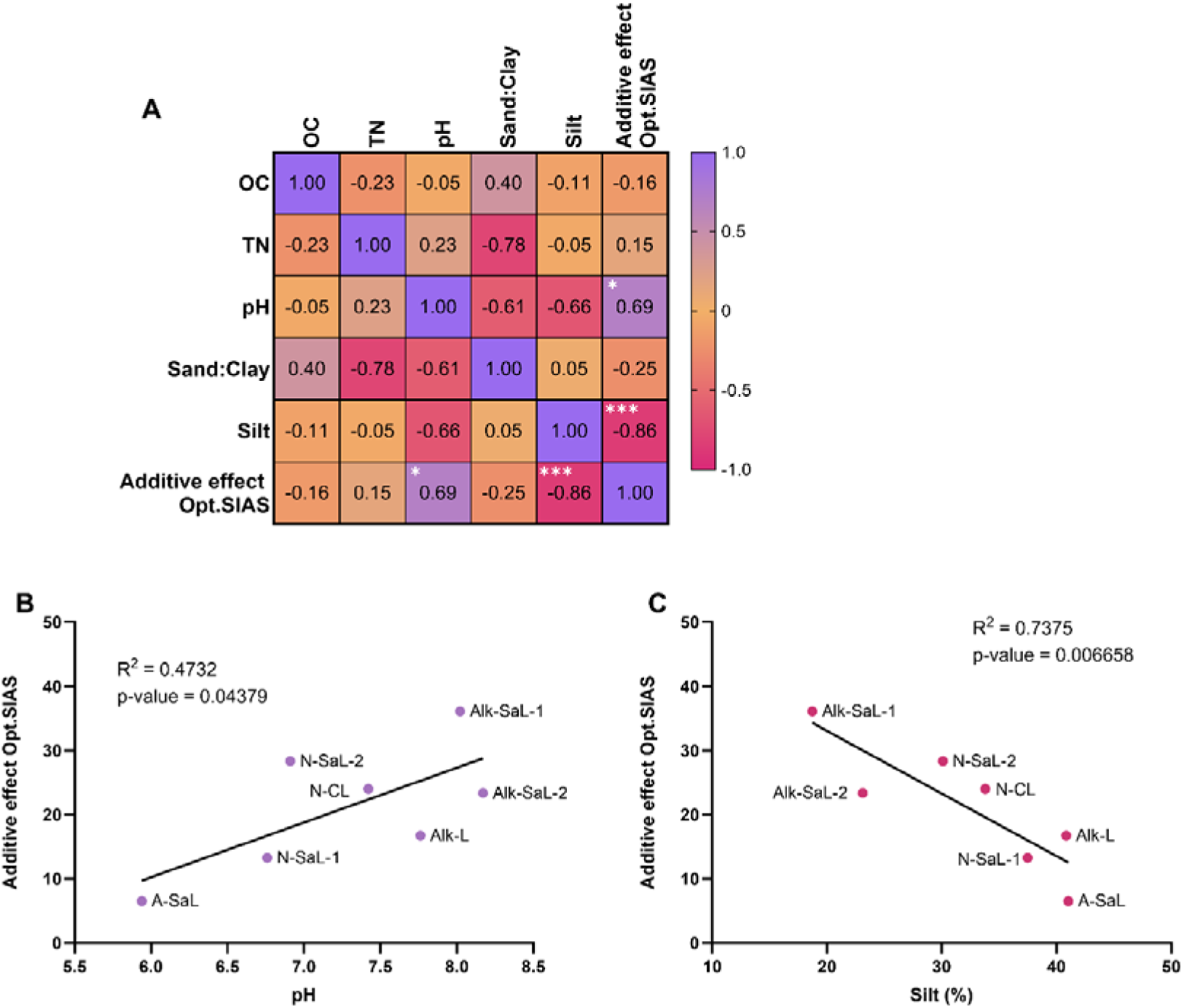
**(A)** Pearson correlations between soil characteristics and the additive effect of Opt. SIAS over DMP. Statistical analysis was assessed via a two-way ANOVA followed by a Dunnett post-hoc test. **(B)** Simple linear regression between soil pH and the additive effect of Opt. SIAS over DMP. **(C)** Simple linear regression between silt content (%) and the additive effect of Opt. SIAS over DMP. OC: organic carbon. TN: total nitrogen. Correlations with an asterisk are significantly different (*: p<0.05; **: p<0.01; ***: p<0.001; ****: p<0.0001).

## 4. DISCUSSION

The identification of Opt. SIAS represents a significant step forward in the development of AOA-targeting NIs. Despite the growing recognition of AOA as important contributors to soil nitrification (Gubry-Rangin et al., 2010; Wang et al., 2021), only a limited number of compounds have been reported to selectively inhibit these microorganisms under soil conditions. Using a soil-based SAR approach, we identified Opt. SIAS as a compound that consistently enhanced nitrification inhibition when combined with DMP across diverse agricultural soils. In contrast to conventional SAR studies of nitrification inhibitors, which are typically performed in pure cultures, our recently developed miniaturized soil incubation assay (Antoniou et al., 2026) enabled the direct evaluation of inhibitor efficacy under environmentally relevant soil conditions and at a throughput sufficient to screen a large compound library. This aspect is particularly relevant given that inhibitor performance in soil cannot always be predicted from pure-culture assays, as previously demonstrated for SIAS-like compounds (Beeckman et al., 2023).

Compared to SIAS, Opt. SIAS contains a carboxamide group at position 4 and two isopropyl substituents at positions 1 and 6, which appear to be key determinants of its enhanced activity in soil. The improved efficacy of Opt. SIAS may be related to changes in lipophilicity. As previously suggested for other NIs, optimal activity likely depends on a balance between sorption to soil particles and mobility in the soil solution (Sidhu et al., 2025; Taggert et al., 2021). Highly lipophilic compounds may become strongly adsorbed onto soil organic matter, reducing their mobility and accessibility to target microorganisms, whereas highly hydrophilic compounds may remain predominantly in the aqueous phase and interact less efficiently with ammonia-oxidizers associated with soil particles. Because AOA can be mobile, but also attach to surfaces such as soil particles (Kerou et al., 2016), successful inhibition likely requires sufficient mobility while maintaining access to particle-associated populations. Opt. SIAS combines two relatively polar moieties (the pyrazolo-pyridine scaffold and the carboxamide group) with several hydrophobic substituents located at the periphery of the molecule. We hypothesize that this combination of polar and non-polar substituents in Opt. SIAS provide a favourable balance between soil mobility, sorption and bioavailability, contributing to its enhanced and more consistent performance across soils.

Although Opt. SIAS displayed robust activity across soils, its efficacy remained influenced by soil physicochemical characteristics. Soil pH and silt content were significantly correlated with the additive effect of Opt. SIAS over DMP. Although both relationships were statistically significant, their interpretation should be approached with caution given the limited number of soils included in the analysis. Still, the strong negative correlation between silt content supports the hypothesis that sorption-related processes influence compound efficacy. Soil texture affects the retention, mobility and accessibility of agrochemicals through interactions with mineral and organic surfaces (Strawn, 2021; Zheng et al., 2025). Consequently, increasing silt content may reduce the bioavailability of Opt. SIAS to target microorganisms, thereby limiting its inhibitory effect. The positive relationship with soil pH likely reflects a combination of physicochemical and biological mechanisms. While pH can influence the behaviour of organic compounds in soil, it is also a major determinant of ammonia-oxidizer ecology and nitrification dynamics (Elrys et al., 2021; Gubry-Rangin et al., 2011). Under alkaline conditions, AOB are generally expected to contribute more strongly to nitrification, while AOA are considered to be more important in acidic soils (Che et al., 2015; Lei et al., 2022). Therefore, at first sight, the positive relationship between soil pH and the additive effect of Opt. SIAS seems to be counterintuitive. However, the additive effect measured here reflects the additional inhibition achieved on top of DMP-mediated AOB inhibition. Under alkaline conditions, nitrification is generally more strongly driven by AOB. Consequently, inhibition of AOB by DMP may increase the relative contribution of residual AOA-driven nitrification (Fan et al., 2022, 2019; Yang et al., 2021; Yin et al., 2021; Zhou et al., 2020), creating a greater scope for additional inhibition by Opt. SIAS. In contrast, in the acidic soil used in this study, application of SIAS alone resulted in stronger nitrification inhibition than in the alkaline soils, consistent with a greater contribution of AOA to nitrification under acidic conditions. However, neither SIAS nor Opt. SIAS enhanced nitrification inhibition when combined with DMP in this soil. This observation highlights an important limitation of inferring the relative contributions of AOA and AOB from inhibitor-based studies. While nitrification inhibitors are frequently used to estimate the importance of different ammonia-oxidizing groups (e.g. Hink et al., 2017; Taylor et al., 2013, 2012), inhibitor responses may not solely reflect the contribution of nitrifiers in untreated soil, but also the community structure that emerges following selective inhibition. This interpretation is further supported by a recent study combining stable-isotope tracing with multiple inhibitor approaches, which demonstrated that the contribution of individual nitrifying guilds cannot always be inferred independently because interactions between groups influence overall nitrification activity (Song et al., 2026). Nevertheless, interpretations based on the classical niche differentiation between AOA and AOB should be made with caution. Although AOA are frequently associated with acidic soils and AOB with neutral to alkaline soils, some exceptions have been reported: some AOB can also grow at very low pH values, whereas several AOA strains can tolerate elevated NH_3_ concentrations, and may remain important contributors to nitrification under alkaline soil conditions (Aigle et al., 2020; Beeckman et al., 2024). Despite this complexity, the additive effect observed when combining Opt. SIAS with DMP further highlights the contribution of AOA to nitrification in agricultural soils, supporting previous observations by Beeckman et al. (2023). Although currently available commercial nitrification inhibitors primarily target AOB, increasing evidence suggests that AOA can significantly contribute to ammonia oxidation under a wide range of soil conditions (Beeckman et al., 2024). The enhanced inhibition obtained through simultaneous targeting of AOA and AOB indicate that effective nitrification management may benefit from the suppression of both functional groups. Interestingly, unlike simvastatin or PTIO, which did not show a measurable additive effect when combined with DMP in soil (Beeckman et al., 2023), Opt. SIAS consistently enhanced nitrification inhibition across multiple soils, highlighting its potential to expand the limited portfolio of soil-effective AOA-targeting nitrification inhibitors.

The identification of Opt. SIAS represents an important step toward the development of effective AOA-targeting nitrification inhibitors. Beyond its value as a complementary research tool for investigating the role of AOA in soil nitrification, Opt. SIAS may also contribute to more effective nitrification management in agricultural systems, potentially reducing nitrogen losses and improving the sustainability of nitrogen fertilization practices. To realize this potential, future studies should evaluate its performance under more agronomically relevant conditions to determine whether the enhanced nitrification inhibition observed here translates into improved nitrogen use efficiency and agronomic benefits. In parallel, comprehensive assessments of its environmental fate and ecotoxicological profile will be required to assess its suitability for agricultural use.

## 5. CONCLUSION

This study demonstrates that soil-based structure-activity relationship (SAR) screening can successfully guide the development of effective AOA-targeting nitrification inhibitors. Using this approach, we identified Opt. SIAS, an AOA-targeting nitrification inhibitor with stronger and more consistent soil efficacy than the original lead compound. The strong additive effects observed when Opt. SIAS was combined with DMP demonstrate the potential of simultaneously targeting AOA and AOB to improve nitrification inhibition and highlight the contribution of AOA to agricultural nitrification, particularly following AOB inhibition. These findings further show that nitrifier contributions inferred from inhibitor studies should be interpreted with caution, as selective inhibition can change the relative importance of non-targeted populations. Collectively, this work expands the limited portfolio of effective AOA-targeting nitrification inhibitors, establishes soil-based screening as a powerful strategy for their optimization, and lays the foundation for next-generation nitrification inhibition strategies aimed at improving nitrogen-use efficiency and reducing the environmental footprint of agricultural fertilization.

## Acknowledgements

This work was supported by the Flemish Agency for Innovation and Entrepreneurship (VLAIO – grant number HBC.2025.1414) and EuroChem Antwerp. We thank Thorsten Wendt and Nils Peters for their input and constructive comments on this work.

## Author contributions

**Conceptualization:** H.M., T.B.; **Methodology:** L.A., F.B., K.V.; **Investigation:** L.A.; **Formal analysis:** L.A., F.B., H.M.; **Supervision:** F.B., T.B., H.M.; **Writing – original draft:** L.A.; **Writing – review & editing:** L.A., F.B., T.B., H.M.

---

**Table S1:**
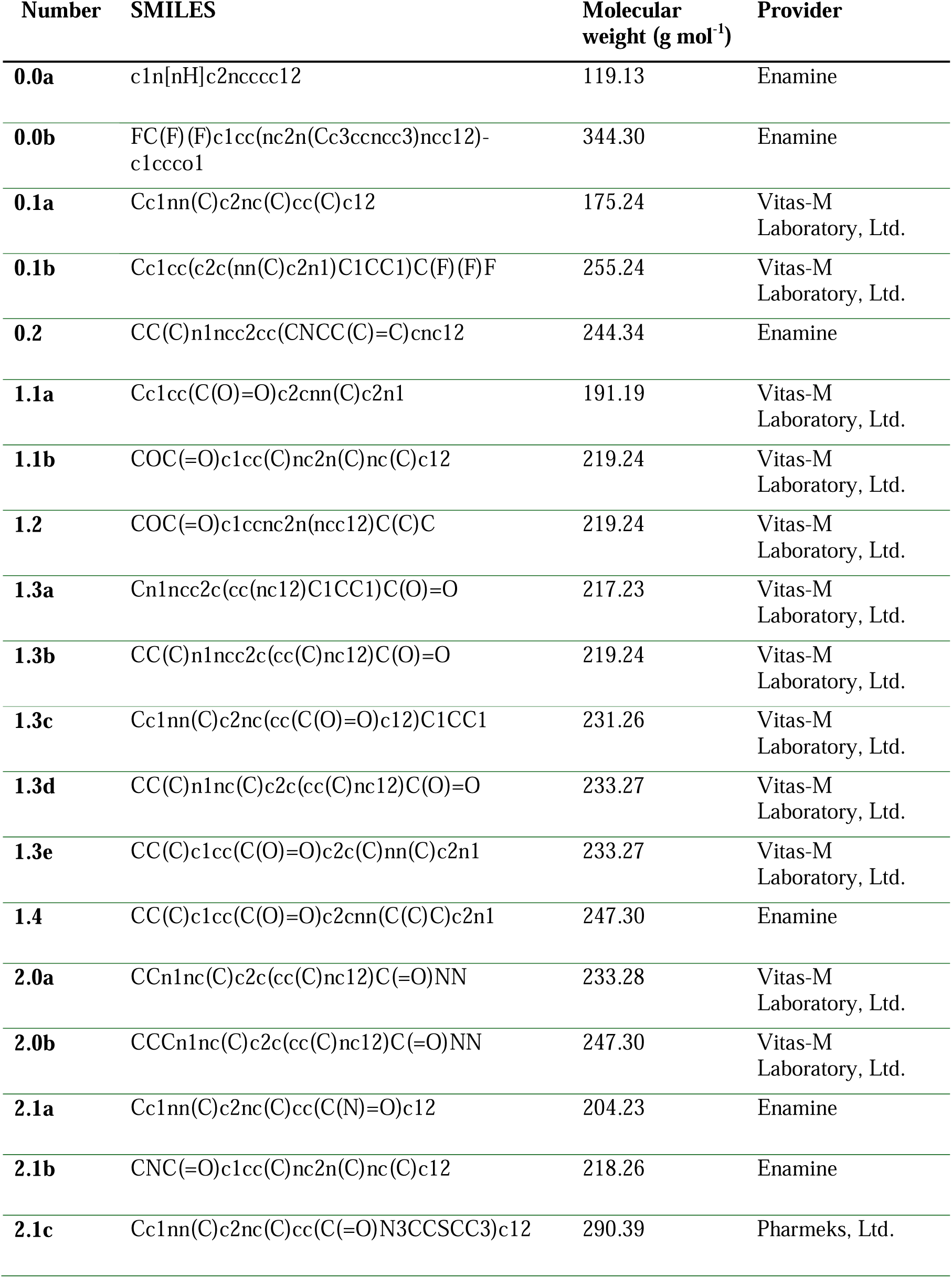
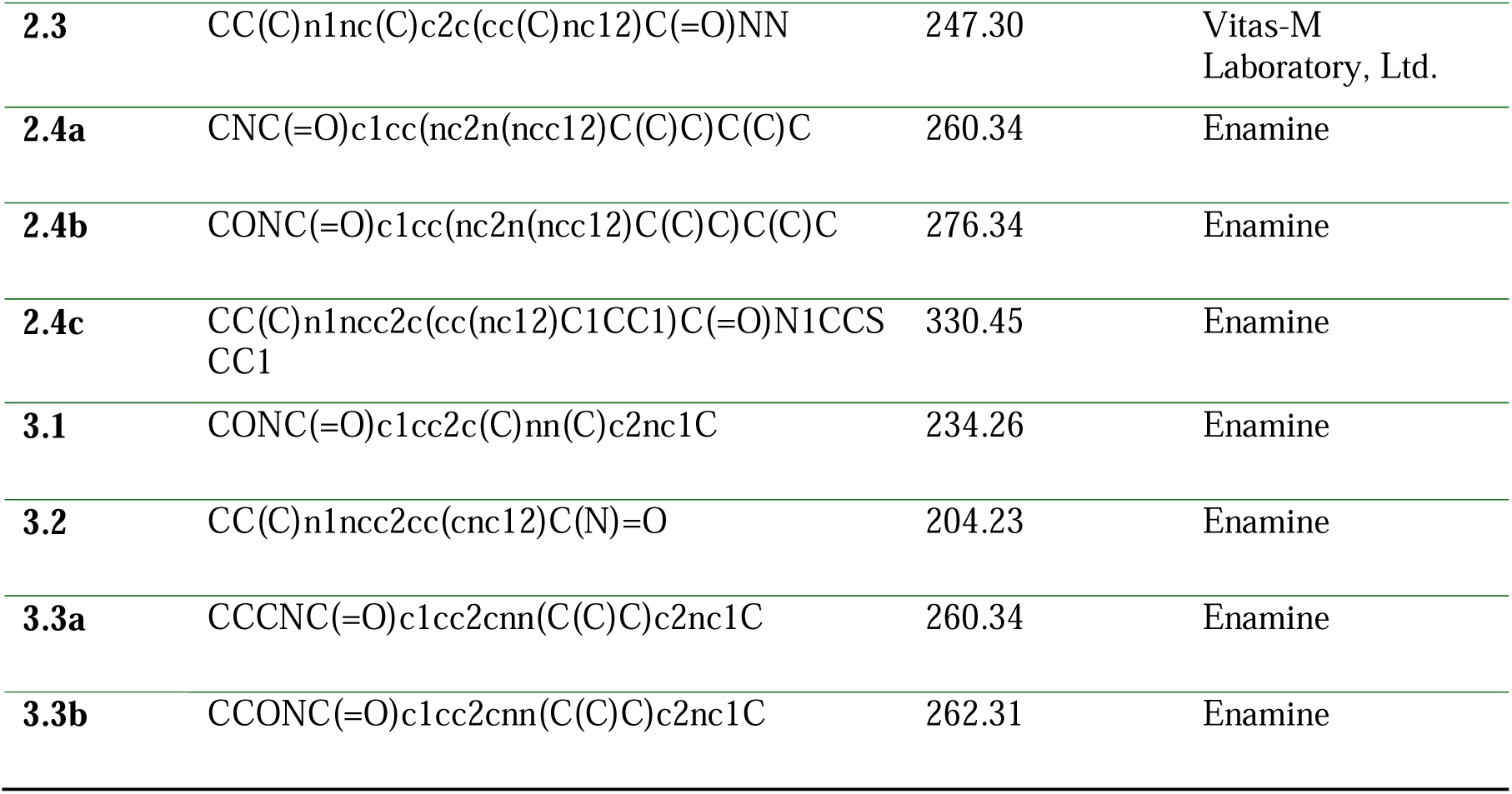
List of the 27 SIAS variants used in the structure-activity relationship (SAR). SMILES: Simplified Molecular Input Line Entry System.

